# Secretory protein Rv1987, a ‘probable chitinase’ from *Mycobacterium tuberculosis* is a novel chitin and cellulose binding protein lacking enzymatic function

**DOI:** 10.1101/2023.08.13.553142

**Authors:** Chiranth M. Prakash, Vani Janakiraman

## Abstract

Bacterial chitinases serve to hydrolyse chitin as food source or as defence mechanism. Given that chitin is not produced by mammals, it is intriguing that *Mycobacterium tuberculosis*, an exclusively human pathogen harbours Rv1987, a probable chitinase and secretes it. Interestingly genes annotated as chitinases are widely distributed among *Mycobacterium tuberculosis* complex species, clinical isolates and other human pathogens *M. abscessus* and *M. ulcerans*. However, Mycobacterial chitinases are not characterized and hence the functions remain unknown. In the present study, we show that Rv1987 is a chitin and cellulose binding protein lacking enzymatic activity in contrary to its current annotation. Further, we show Rv1987 has moon lighting functions in *M. tuberculosis* pathobiology signifying roles of bacterial cellulose binding clusters in infections.

## Introduction

*M. tuberculosis* (Mtb), the causative agent of tuberculosis, is one of the oldest known human pathogens and still causing robust disease that is responsible for over 2 million deaths annually [1]. Mtb also causes latent infections that can get reactivated anytime during one’s life time leading to serious clinical consequences [2]. Mycobacteria are intra-cellular pathogens that have developed specific mechanisms to invade and survive within their host making the host-pathogen interplay in tuberculosis infection is very complex. Therefore, examining the mechanisms by which, *M. tuberculosis* activates or evades the host responses is a key to developing a greater understanding of the pathology of tuberculosis. In this context, we report characterization of Rv1987, a possible chitinase from *M. tuberculosis*.

Bacterial chitinases have always been viewed as enzymes that either hydrolyse chitin as a food source especially in environmental bacteria or that serve as a defence mechanism against organisms such as fungi that contain chitin as a major structural component. Given the fact that chitin is not produced by mammals, it is intriguing that *M. tuberculosis*, which is an exclusively human pathogen, harbours the gene Rv1987 coding for probable chitinase. Recent evidences show that the functions of microbial chitinases are not limited to their enzymatic activity and that chitinases are associated with virulence [3].

As chitin is one of the most abundant polysaccharides on earth, second only to cellulose, and is found primarily in fungi and the exoskeletons of arthropods, studies on polysaccharide utilizing locus (PULs) in bacteria including gut bacteria is gaining interest [4]. Interestingly, genes annotated as chitinases are widely distributed among *Mycobacterium tuberculosis* complex (MTBC) that cause tuberculosis in different organisms and in non-tuberculous, human pathogenic mycobacteria such as *M. abscessus* and *M. ulcerans*. However, Mycobacterial chitinases are not characterized and the purpose of their presence in Mycobacterial genome remains enigmatic. Therefore, we studied Rv1987, a possible chitinase from *M. tuberculosis* to understand its features and role in Mycobacterial physiology.

Rv1987 is a secreted protein and is part of RD2 locus in *M. tuberculosis* [5]. Regions of differences (RDs) are sequences that are absent in the attenuated vaccine strain, *M.bovis* BCG in comparison to the virulent strain *M. tuberculosis* H37Rv. Proteins encoded within these regions are thought to be contributing for virulence and deletion of these regions could lead to attenuation of *M. bovis* BCG [5]. This also hints towards the potential role of *M. tuberculosis* encoded protein product of Rv1987 as a possible virulence factor. This protein has also been explored as a possible vaccine candidate [6]. In the present work, we have characterized Rv1987 and show that it is a novel chitin and cellulose binding protein of Mycobacteria lacking enzymatic activity in contrary to its current annotation and has moon lighting functions.

## Materials and Methods

### Cloning, expression and purification

Plasmid harbouring Rv1987 gene was amplified by PCR using gene specific primers. This master clone pETG28 was received as a kind gift from Dr.Bernard Henrissat of AFMB, CNRS-Aix-Marseille University, France. The recombinant construct was transformed and into *E.coli* BL21 DE3 expression system. The recombinant protein was expressed by inducing the expression using 0.5 mM IPTG and was incubated at 21°C for 16 hours. Post induction with IPTG, protein was found to be present both in the pellet and supernatant fractions. Since majority of the protein was found to be in the pellet fraction, the protein was solubilized with a mild detergent N-lauryl sarcosine (NLS). Protein was then purified using Ni-NTA agarose column using standard procedures. Purified protein product was confirmed through 12% SDS-PAGE gel and subsequent western blot using anti-His antibodies. Protein concentration throughout the study was determined *via* Bradford assay (catalogue number-VWR M172).

### Analysis of catalytic activity

The ability of purified protein Rv1987 to hydrolyse polymeric carbohydrates (chitin and cellulose) was determined by 3,5-di-nitrosalicylic acid (DNS) method a well-established method [7–9]. Different concentrations of purified Rv1987 was incubated with colloidal chitin (from the shrimp shell-HiMedia) and carboxymethylcellulose (HiMedia). Briefly, 0.1% colloidal chitin in sodium acetate buffer (pH 5.0) and 0.1% carboxymethyl cellulose in citrate buffer (pH 5.0) were used to access the chitinase and cellulase activity respectively. Purified protein Rv1987 and substrate(s) were incubated for 30 minutes at 50°C. Different concentration ranges of Rv1987 was 5,10, 20, 50 ug/ml while 5 ug/ml and 10 ug/ml for cellulase standard (from *Tricoderma reseei*) were used. Equal volumes of freshly prepared DNS reagent (consisting of 1% di-nitrosalicylic acid, 0.2% phenol, 0.05% sodium sulfite, and 1% sodium hydroxide) was added, followed by boiling in a water bath for 5 minutes. Samples were centrifuged and supernatant was collected and absorbance of the reduced sugars was measured at 540 nm. To estimate the total reducing sugars, obtained values were compared to a standard curve plotted using N-acetyl glucose amine and D-glucose for chitinase and cellulase respectively.

### Analysis of carbohydrate binding properties

Owing to the absence of catalytic activity in the previous experiment, we checked if the protein exhibits any carbohydrate binding property. Firstly, ability of Rv1987 to bind to avicel (cellulose substrate) was analyzed using cellulose binding assay. Briefly, 8 mg/ml of substrate was washed and equilibrated with 20 mM sodium citrate buffer. 6 µg/ml of purified Rv1987 was transferred to the equilibrated avicel. This suspension was incubated for 4 hours at room temperature with gentle mixing. Supernatant was collected after centrifugation, while pellet was washed with citrate buffer to remove any unbound protein. The bound and unbound proteins were quantified using Bradford assay and the difference in the concentration revealed that Rv1987 was bound to chitosan. In parallel, the samples were run and visualized on 10% SDS-PAGE.

Similarly, ability of Rv1987 protein to bind to colloidal chitin from shrimp and chitosan (chitin substrate) was confirmed using chitin binding assay. The substrate (8 mg/ml) was washed and equilibrated with 50 mM sodium phosphate (pH 6.5). 6 µg/ml of purified Rv1987 was transferred to the equilibrated substrate (colloidal chitin and chitosan). The suspension was incubated for 24 hours at room temperature with gentle mixing. The supernatant was collected after centrifugation, while pellet was washed with sodium phosphate buffer to remove any unbound protein. The presence of bound protein was analysed by addition of 10 µl of 6x SDS gel loading dye to substrate. Bound and unbound proteins were quantified using Bradford assay and visualized on 10% SDS-PAGE.

### Sequence and domain organization

Amino acid Sequence of Rv1987 from *M. tuberculosis* H37Rv, the routinely used virulent strain was retrieved (Uniprot:P9WLQ1.1). Analysis for presence of functional domains was performed using conserved domain database (CDD) available at NCBI[10].

### Sequence based analysis with glycoside hydrolases

Amino acid sequences of the glycoside hydrolases (GH18 family) was retrieved from UniProt [11]. These sequences were aligned with Rv1987 sequence using online multiple sequence alignment tool praline (http://www.ibi.vu.nl/programs/praline/)[12]. Default parameters were applied and the aligned sequences were inspected for the number of gaps and insertions. In order to further scrutinize, sequences of other known bacterial chitinases, bacterial chitin binding proteins were also retrieved and multiple sequence alignment was performed as mentioned above.

### Structure based analysis

As the sequence-based studies are only indicative, further analyses was carried out employing structure-based analyses. The modelled structure of Rv1987 was retrieved from Mycobacterial structural genomics database (http://proline.biochem.iisc.ernet.in/sincre/). To assess and validate the stereo chemical quality and structural integrity of the model, an online tool RAMPAGE (http://mordred.bioc.cam.ac.uk/~rapper/rampage.php) was used. To define the equivalence map between residues for all the chosen structures based on their relative position in space, structure-based alignment was performed. We had to resort to this structure due to the non-availability of experimentally solved crystal structure. However, we have made all efforts towards the quality assessment of the structure. The database also shows more than 75% of model coverage for the Rv1987 with the template structure.

All available experimentally solved structures of known bacterial chitinases, cellulases, chitin binding proteins and cellulose binding proteins were retrieved from PDB and alignment was performed. PyMOL script was used for the alignment and root mean square deviation (RMSD) of the superimposed structures was calculated and analysed.

### Analysis of binding pockets of Rv1987 and substrate specificity

To gain insights into binding pockets and plausible substrates recognized by Rv1987, docking analysis was performed. To evaluate the substrate preference, 14 different compounds were selected from Pubchem [13] based on similarity in chemical nature and molecular weight. Retrieved 2D formats of the substrates were then converted to 3D formats using Open Babel [14]. All 14 chosen substrates were analysed for binding with Rv1987 using docking analysis performed using AutoDock vina [15]. Grid maps were generated using AutoGrid program in AutoDock4.2. Grid box was made up of 72 × 52 × 48 grid points in x, y and z dimensions respectively with a spacing of 0.375 Å to cover the entire protein-substrate. Substrates were assigned as ‘flexible’ with the number of rotatable bonds set as 8/32. Docked configurations were then analysed for all the interactions between the substrates and the protein using Protein-Ligand Interaction Profiler tool [16].

### Analysis of subcellular localization of Rv1987

Several Mycobacterial proteins, in addition to being secreted into culture supernatants have been shown to be retained in the cell wall/membrane performing modulatory functions at such peripheral structures or adhesion functions [17]. The current functional classification of mycobacterial proteins predicts Rv1987 under cell wall and cell processes [18]. Therefore, to assess the nature of sub-cellular localization of Rv1987 in *M. tuberculosis* which in turn will help to establish its further functional role, localization studies were performed using different cellular fractions from the virulent strain *Mtb* H37Rv procured from BEI (Biodefense and Emerging Infections Research Resources Repository) resources [19]. Custom made rabbit polyclonal antibodies against purified Rv1987 were generated and procured from Vipragen, Mysore, India. Briefly, different cellular fractions such as total soluble cell wall proteins, culture filtrate fraction, cell membrane fraction and whole cell lysates were resolved by SDS-PAGE and western blotting was performed using anti-rabbit polyclonal antibody raised against Rv1987 as primary antibody at 1:5000 dilution. The blot was developed using goat anti-rabbit HRP antibody and visualized.

### Generation of recombinant M. smegmatis expressing Rv1987

Rv1987 gene was also cloned in the Mycobacterial shuttle vector pVV16 for over expression in *M. smegmatis* to assess its functional role in Mycobacterial physiology. Briefly, the shuttle vector pVV16 and recombinant clone pVV16::Rv1987 were electroporated into freshly prepared electrocompetent cells. Recombinant clones were selected by plating on 7H10 agar plates containing kanamycin (50 µg/ml) and hygromycin (100 µg/ml) as the selection markers. Expression of Rv1987 in the recombinant clones was confirmed through western blot using specific antibodies.

### Interaction of human alveolar epithelial cells and monocytes with Rv1987

Interaction of Rv1987 with human cells was assessed using human alveolar type-II pneumocyte cell line A549 and human monocyte cell line THP-1 respectively to assess its role in adhesion and in rendering biological fitness in terms of intracellular survival of bacteria. For cytoadherence assay, A549 cells were seeded at a density of 1×10^5^ cells/well in 35 mm dishes. Cells were allowed to adhere. Recombinant *M. smegmatis* (over expressing Rv1987) culture pellet from log phase cultures was washed and used for infecting cells at an MOI (multiplicity of infection (MOI) of 10. Cells and bacteria were incubated for 60 minutes. Cells were washed with PBS and treated with 100 µg/ml of gentamycin to remove any residual extracellular bacteria. Cells were then lysed and the whole cell lysates were diluted and plated. Similarly, to study intracellular survival, THP-1 cells were seeded and differentiated with 20 ng/ml PMA (phorbol 12-myristate 13-acetate). Subsequently, cells were washed and the differentiated THP-1 cells were seeded at a density of 1×10^5^ cells/well in 35 mm dishes. Cells were then infected with an MOI of 10 for 24, 48 and 72 hours and assessed for the viability of intra cellular bacteria on preparing lysates (using cell lysis buffer (0.1% triton-X 100 in PBS for 20 minutes. Lysates were then diluted and plated on 7H10 media (media for growth of *M.smegmatis*) plates containing kanamycin (50 µg/ml) and hygromycin (100 µg/ml) as antibiotic selection markers.

## Results

*M. tuberculosis* encoded protein Rv1987 is of 142aa with the region 40-139aa being a conserved domain occupying the majority of the protein. Though it is presently annotated as ‘possible chitinase’ with function denoted as chitin hydrolysis in the whole genome sequence of *M. tuberculosis* [20][21], it is interesting to note that this conserved domain is structurally predicted to be a cellulose binding domain (Figure 1a) belonging to carbohydrate binding module-2 (CBM-2) family. Therefore, we hypothesized that the presence of this domain probably implies that Rv1987 is also a cellulase/cellulose binding protein.

**Figure 1.**
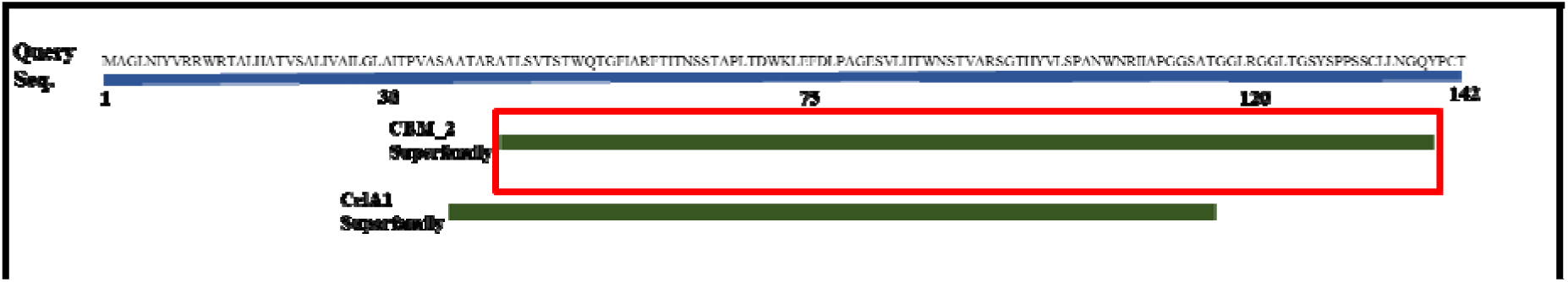
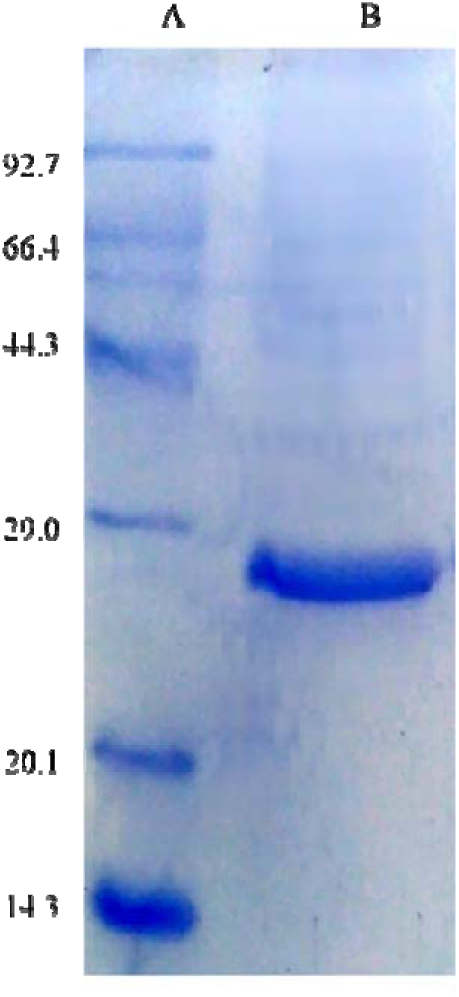
**a. Pictorial representation of conserved domain of Rv1987**. Prediction results from conserved domain database (CDD) search have been adapted and represented. It can be observed that between 40-139aa region is a conserved domain belonging to the carbohydrate binding module-2 superfamily (highlighted in red). CDD also predicted a significant homology to CelA1 domain superfamily (involved in carbohydrate transport and metabolism). **b. Expression and purification of Rv1987.** SDS-PAGE showing expressed and purified recombinant protein Rv1987. Lane A: Molecular weight marker (kDa), Lane B: Purified protein Rv1987 with thioredoxin domain (∼26.1kDa).

Features of carbohydrate-binding modules (CBMs), a contiguous sequence (∼100 aa) found within large number of carbohydrate-active bacterial enzymes and that could act as functional domains have been comprehensively presented in CAZy database [22][23]. These were originally classified as cellulose-binding domains (CBDs) based on the initial discovery of several of these modules that bound cellulose [24][25]. However, with recent observations of modules that bind carbohydrates such as chitin and xylan yet sharing the similar features of CBDs has necessitated the classifications into families [23][26]. Interestingly, independent putative CBMs like Rv1987 are rare occurrences.

### Rv1987 does not hydrolyse chitin/cellulose

*E.coli* BL21-DE3 expressed Rv1987 was purified and was confirmed using anti-His antibodies (Figure 1b). We next checked for catalytic activity of purified Rv1987 against colloidal chitin for its ability to cleave chitin. Results are represented in Figure 2a. There was no liberation of free glucose amine as measured through the DNS assay. This indicated that purified protein Rv1987 failed to demonstrate any chitinase activity, even after incubating with a higher concentration of (50 µg/ml) of purified protein.

**Figure 2.**
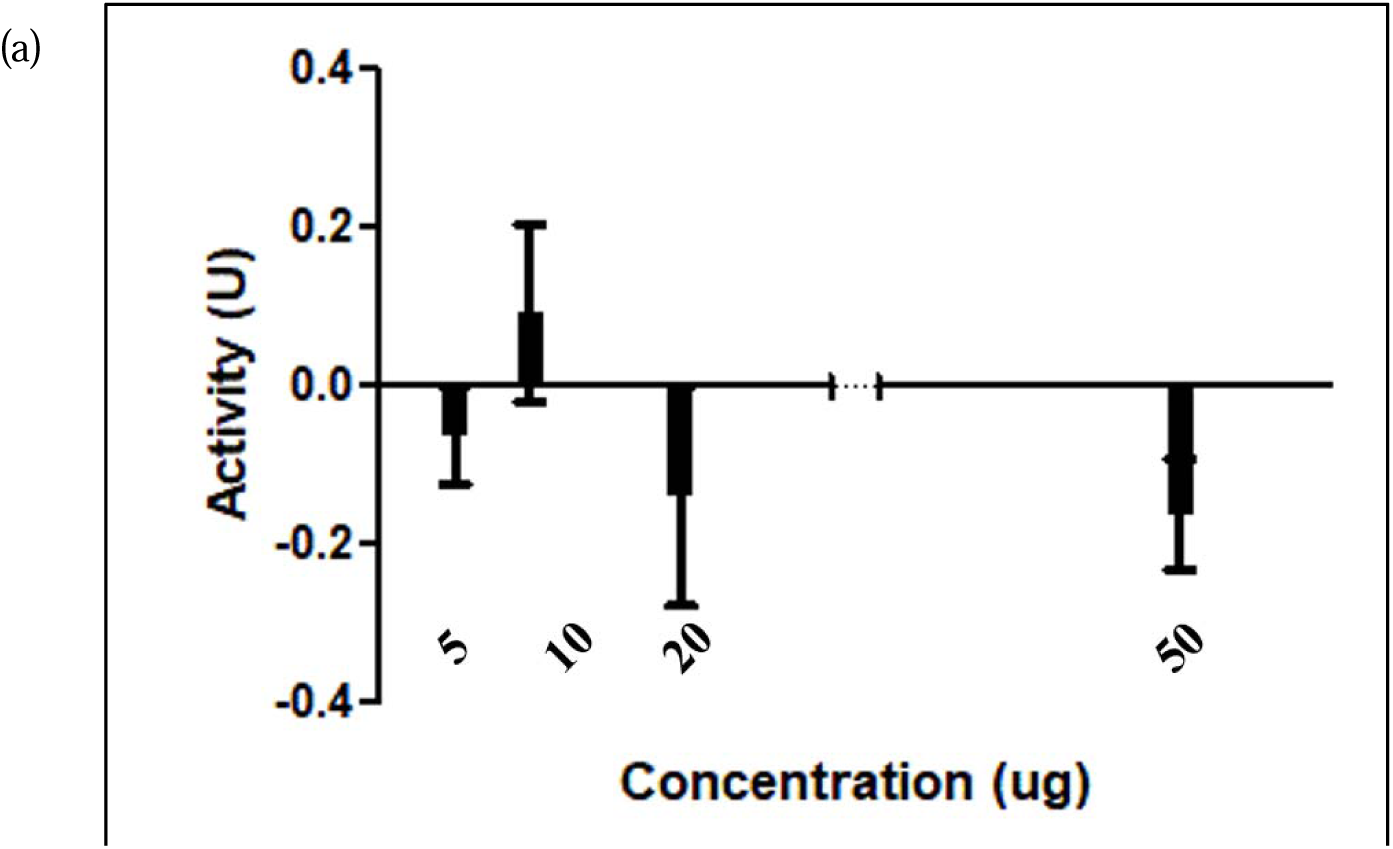
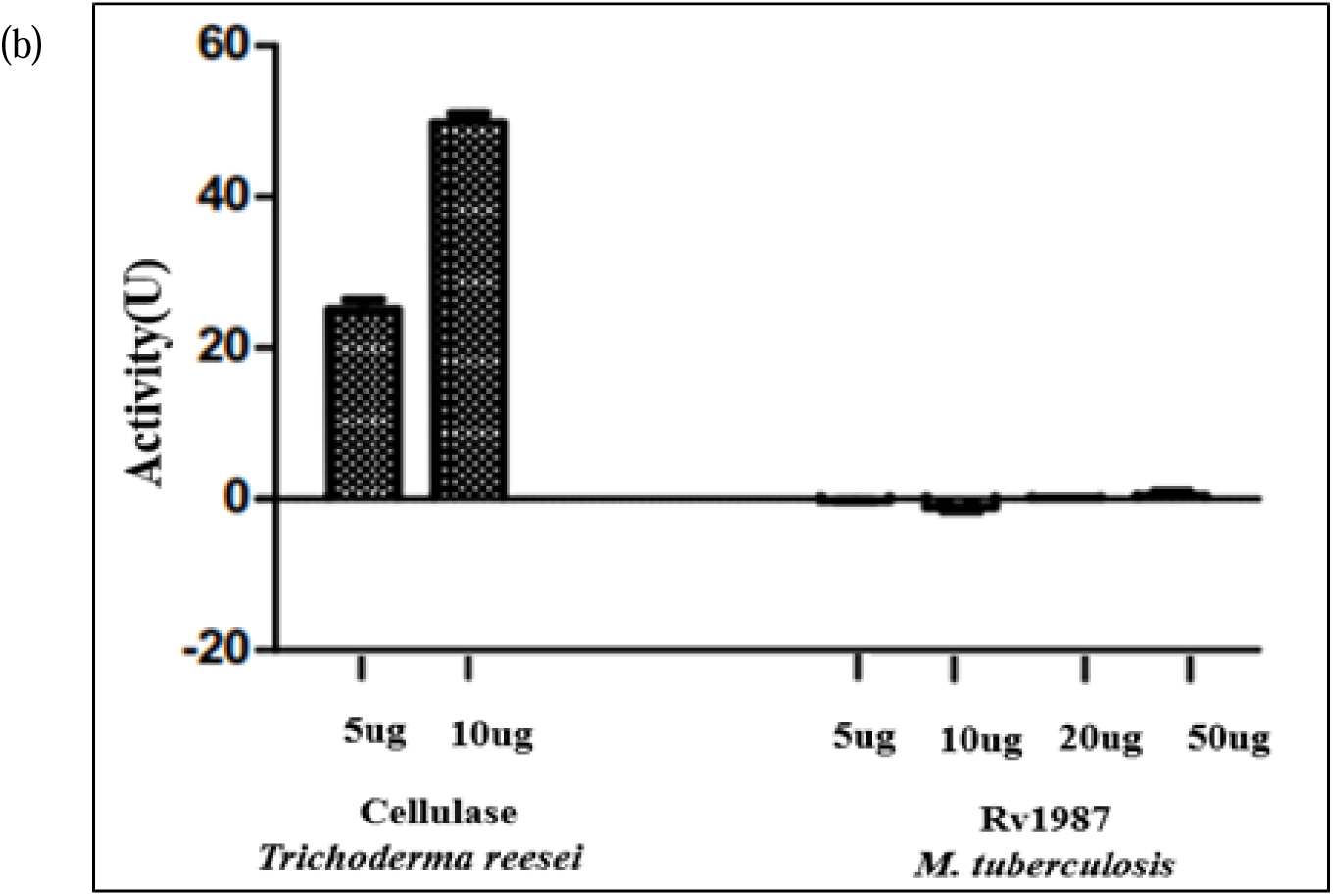
Analysis of catalytic activity of Rv1987. **(a) Analysis of chitinase activity.** Catalytic activity was assessed by 3,5-di-nitrosalicylic acid (DNS) method as described in the methods section. Concentration range of purified Rv1987 used is between 5-50 µg. Absence of chitinase activity indicates the inability of Rv1987 to hydrolyse chitin. **(b) Analysis of cellulase activity.** Catalytic activity was assessed using cellulase from *Trichoderma reesei* which exhibited enzymatic activity and there was no such activity observed with purified Rv1987.

Similarly, assays to assess for cellulase activity were performed. Results in Figure 2b are from cellulase activity assay performed using cellulase from *Trichoderma reesei* and purified Rv1987. It was observed that on treatment of different concentrations (5 µg/ml,10 µg/ml) of enzyme with substrate CMC, the activity was 20 U and 50 U in case of *Trichoderma* cellulase. However, purified Rv1987 did not exhibit any cellulase activity when subjected to a lower to higher concentration range (5 µg/ml to 50 µg/ml). Thus clearly indicating that Rv1987 does not hydrolyse cellulose.

### Rv1987 exhibits binding to chitin/cellulose

We next checked, though there is no catalytic activity, if Rv1987 exhibits binding to these polysaccharide substances. On subjecting purified Rv1987 for interaction with three different substrates including chitin, chitosan and carboxymethylcellulose, we observed prominent a binding. The overall results (figure 3a-3f) indicate that Rv1987 via its carbohydrate binding module (CBL) directly interacts with chitin and cellulose. Also, quantification of different fraction washes through Bradford assay implied the presence of protein bound to substrates. Recent studies have demonstrated that cellulose is the main component of *M. tuberculosis* biofilms [27], [28]. Interestingly, here we have demonstrated an unexpected mode of action of a Mycobacterial carbohydrate binding protein (CBP) that could possibly entail a direct interaction with Mycobacterial biofilms.

**Figure 3.**
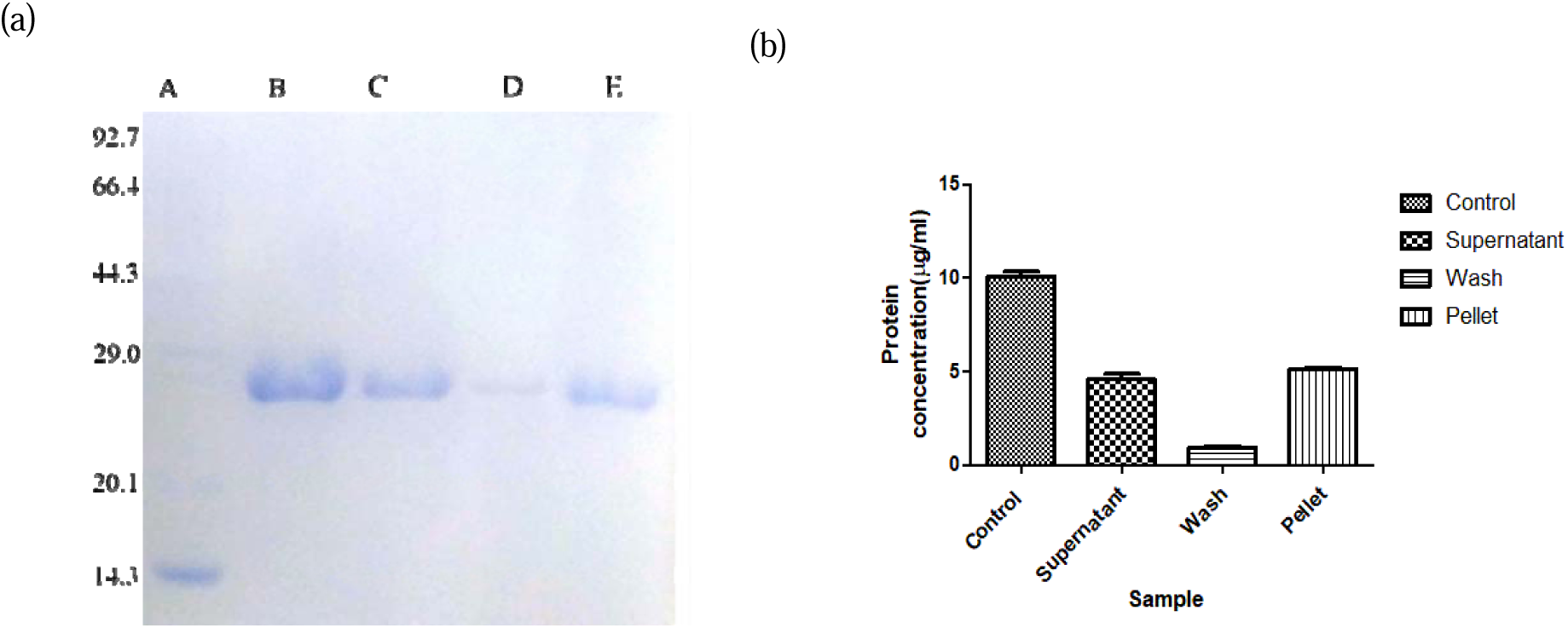
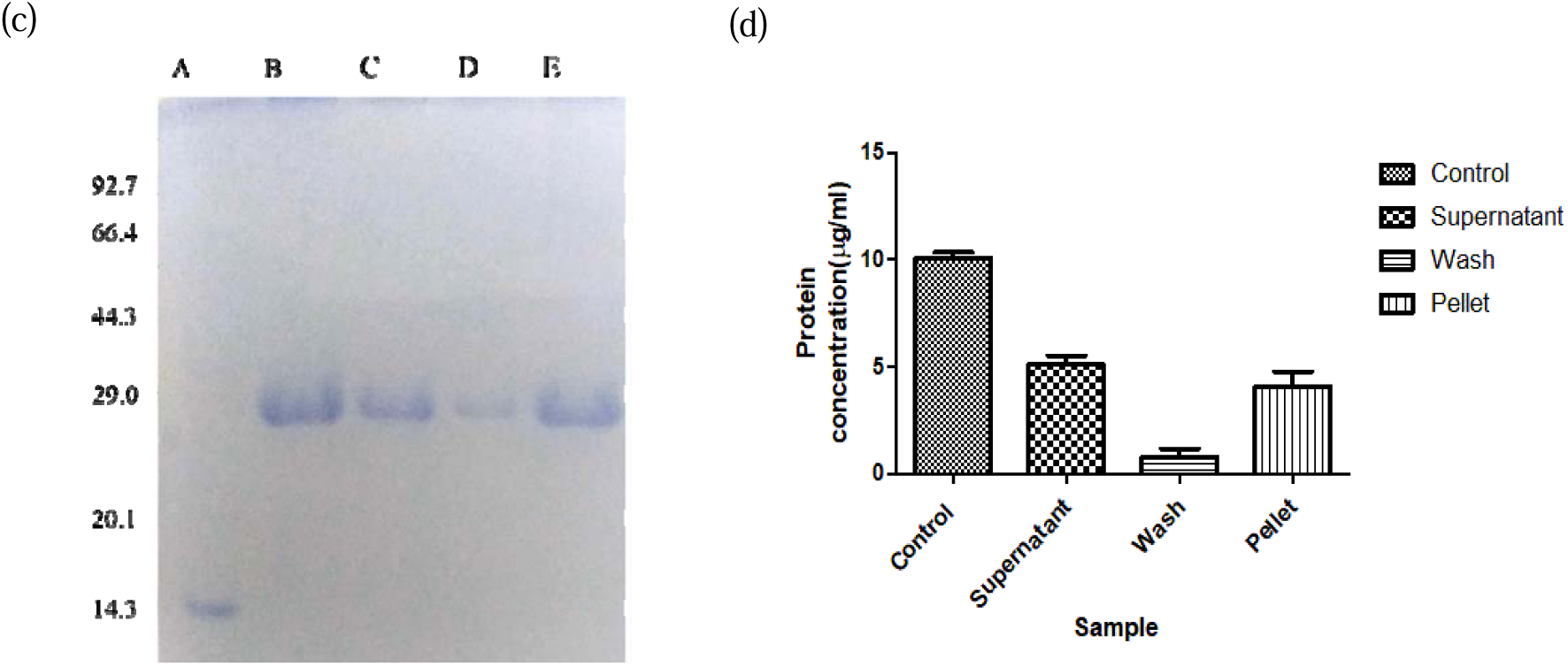
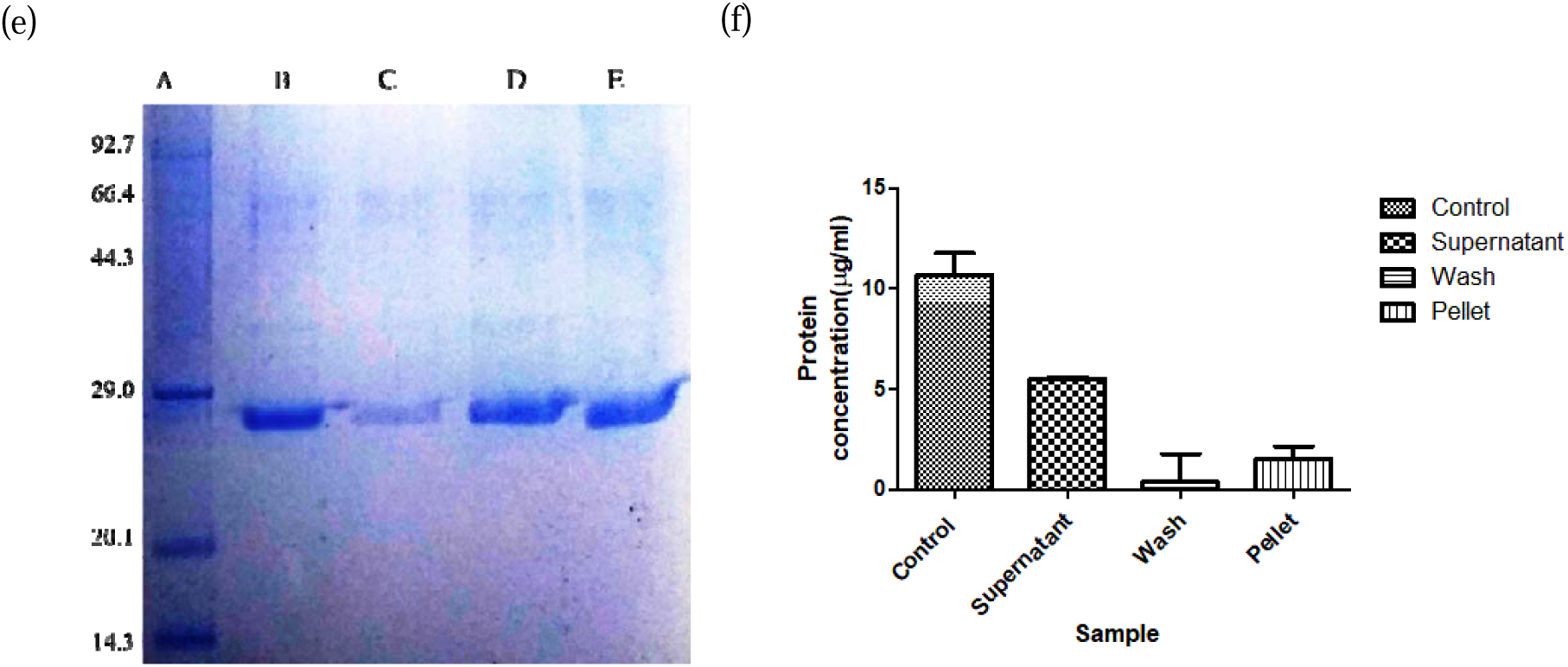
Analysis of carbohydrate binding properties of Rv1987. **(a,b)** Chitin binding assay of Rv1987 visualized on 10% SDS-PAGE (a) Lane A: Marker, Lane B: Rv1987 (6 µg/ml), Lane C: Unbound protein, Lane D: Buffer Wash, Lane E: Bound protein (b) Quantification of protein fractions through Bradford assay. Error bars indicate standard deviation. **(c, d)** Chitosan binding assay of Rv1987 visualised on 10 % SDS-PAGE (c) Lane A: Marker, Lane B: Rv1987 (6 µg/ml), Lane C: Unbound protein, Lane D: Buffer Wash, Lane E: Bound protein (d) Quantification of protein fractions through Bradford assay. Error bars indicate standard deviation. **(e, f)** Cellulose binding assay of Rv1987 visualized on 10% SDS-PAGE (e) Lane A: Marker, Lane B: Rv1987 (6 µg/ml), Lane C: Unbound protein, Lane D: Buffer Wash, Lane E: Bound protein (f) Quantification of protein fractions through Bradford assay. Error bars indicate standard deviation.

### Rv1987 lacks the signature motif of GH18 family hydrolases

A total of 44 sequences belonging to GH18 hydrolase family were retrieved and aligned with the sequence of Rv1987. It was found that GH18 hydrolase family consists of a consensus sequence DGXDXDXE (Figure 4a). The characteristic sequence and structural features of GH-18 family is exhaustively described in the carbohydrate-active enzyme (CAZy) database [22] [23]. Presence of glutamic acid in the consensus region is known to cause protonation of oxygen in scissile glycosidic bonds that results in the hydrolase activity. Strikingly, this signature motif was not present in Rv1987. Absence of this consensus sequence in Rv1987 implies that the protein should not be grouped under the GH18 family as in the present annotation of the whole genome sequence of *M. tuberculosis*.

**Figure 4.**
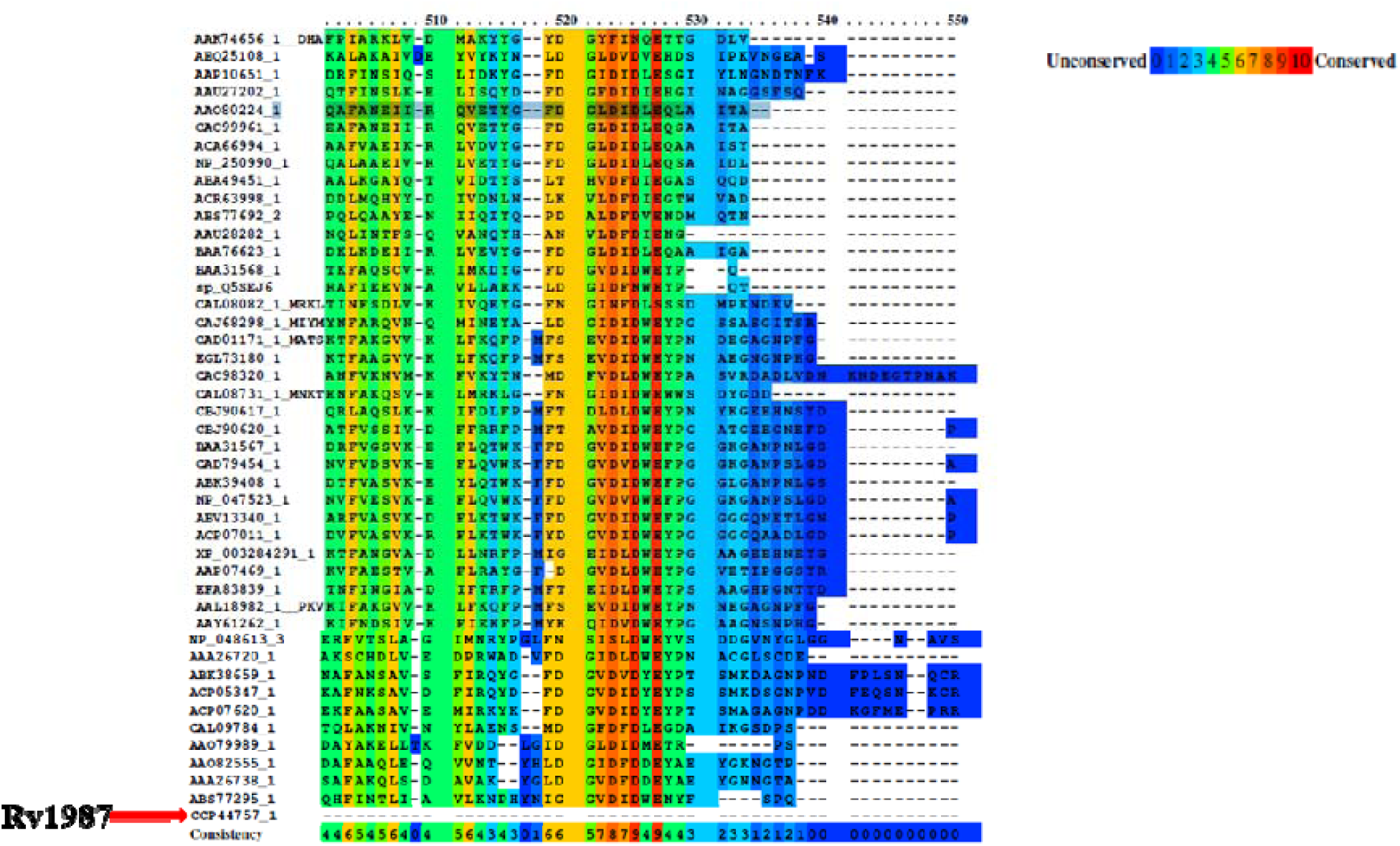
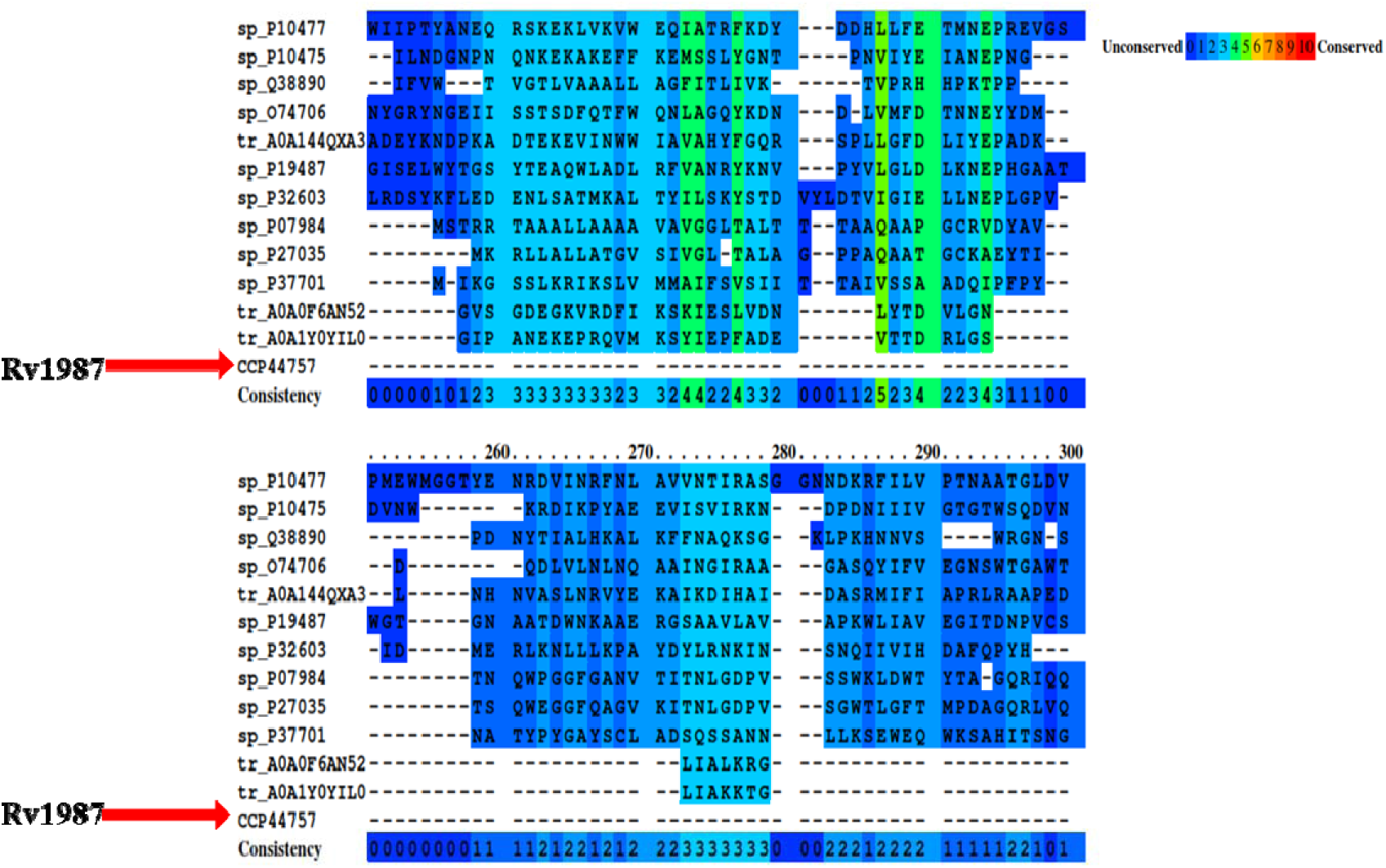
**a. Multiple sequence alignment of GH18 Chitinases with Rv1987.** Absence of signature sequence motif DGXDXDXE in Rv1987 in comparison to all other bacterial chitinase belonging to GH18 family. **b. Multiple sequence alignment of Rv1987 with bacterial cellulases**. Rv1987 does not share any sequence similarity with bacterial cellulases.

Further, sequence alignment of Rv1987 against other bacterial chitinases and bacterial cellulases (Figure 4b and 4c) showed no identical residues in the conserved region of the active site of the catalytic domains. In contrary, several regions of Rv1987 were found to have identical amino acids and shared conserved regions when aligned with bacterial chitin binding proteins (Figure 5a), indicating that Rv1987 could be a chitin binding protein.

**Figure 5.**
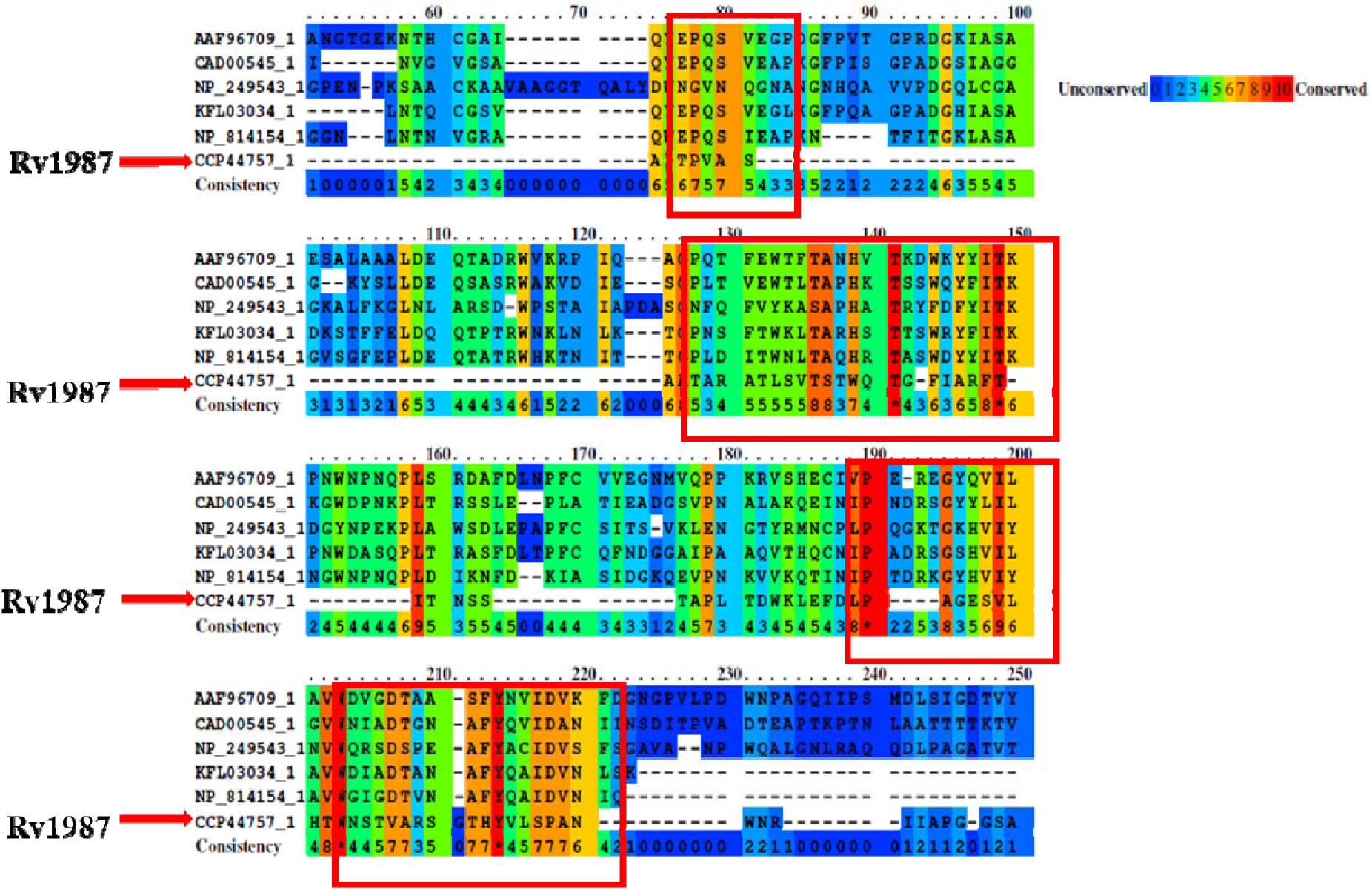
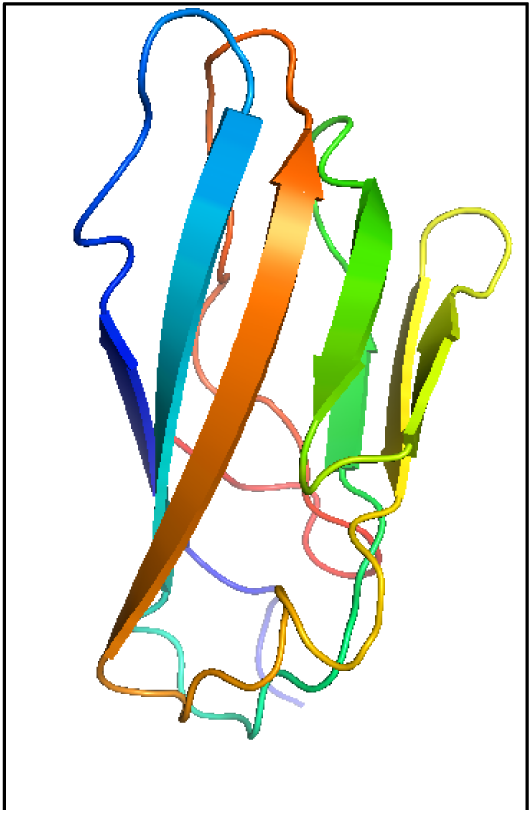
**a. Multiple sequence alignment of Rv1987 with bacterial chitin binding proteins.** Rv1987 shares significant sequence similarity with bacterial chitin binding proteins. **b. Structure of Rv1987 from SInCRe database.** The modelled structure is largely composed of β-sheets and turns. The structure was subjected to quality check as mentioned in the methods section before proceeding with further analyses.

### Rv1987 structurally matches with chitin binding domains of chitinases and not with catalytic domains

The modelled structure of Rv1987 was accessed from the SInCRe database (Figure 5b) and is found to be composed of β-sheets and turns. To assess the stereo chemical quality and structural integrity of the model, RAMPAGE tool was used to generate Ramachandran plot which showed more than 80% amino acids in the favoured regions.

Identification of structurally similar proteins would help in revealing differences and similarities between related molecules thereby allowing inferring of functional properties. Therefore, all relevant proteins (bacterial chitinases, cellulases, chitin binding proteins and cellulose binding proteins) for which crystal structures are available till date were curated. Structural alignment was carried out and the modelled structure of Rv1987 was superimposed with each one of these structures. Root mean square deviation (RMSD) between the structures was considered as the index of alignment accuracy. In protein structure prediction/similarity-based studies, usually RMSD between predicted and experimental structures is considered as a measure to assess the similarity between structures. As a standard, RMSD (< 3 å) is considered as a threshold for scoring as closely homologous proteins [29–31]. Strikingly, structural alignment of Rv1987 with several crystal structures of bacterial chitinases showed no similarity. Structural alignment of Rv1987 with various bacterial cellulases also showed no similarity.

Interestingly, a particular domain of chitinase A from *Vibrio harveyi* (3b9d) which happens to be a chitin binding domain (and not the catalytic domain) had high similarity with Rv1987 (Figure 6). This prompted us to further analyse Rv1987 proteins’ similarity with chitin/cellulose binding proteins. Structural alignment of Rv1987 with crystal structures of chitin binding proteins (Table 1) show that Rv1987 is similar to chitin-binding domain (2rtt) of *Streptomyces coelicolor* with RMSD of 2.066 (Figure 7). But most notably, alignment with cellulose binding domain containing proteins (Table 2) showed that Rv1987 exhibits very least structural deviation with cellulose binding domain of *Cellulomonas fimi* (1exg) cellulase (experimentally solved crystal structure) with a RMSD of 0.289. This close structure resemblance further reiterates the quality of the model structure used for the studies. Also, it is interesting to note that this protein also belongs to CBM-2 family as that of the conserved domain of Rv1987 (Figure 1a).

**Figure 6.**
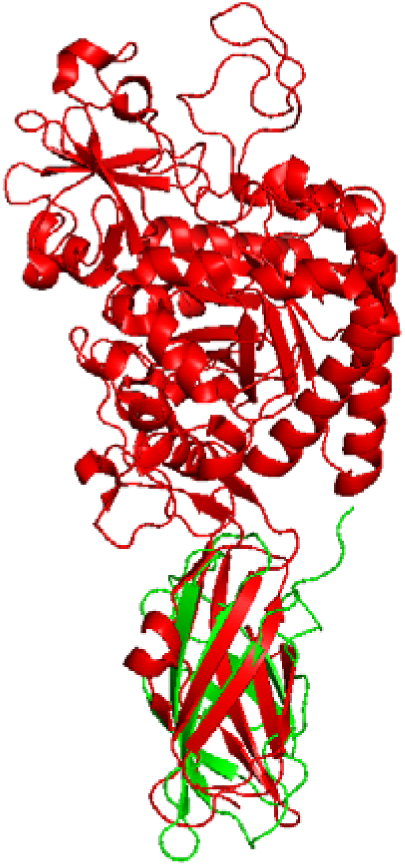
Structure based alignment of Rv1987 with *Vibrio harveyi* chitinase A. Rv1987 was aligned (represented in green color) with a well characterized structure of chitinase A from *Vibrio harveyi* (3b9d) (represented in red color). It is of interest to note that though the full length chitinase was aligned, only a particular domain of chitinase A, which is a chitin binding domain (and not the catalytic domain) has high similarity with Rv1987.

**Figure 7.**
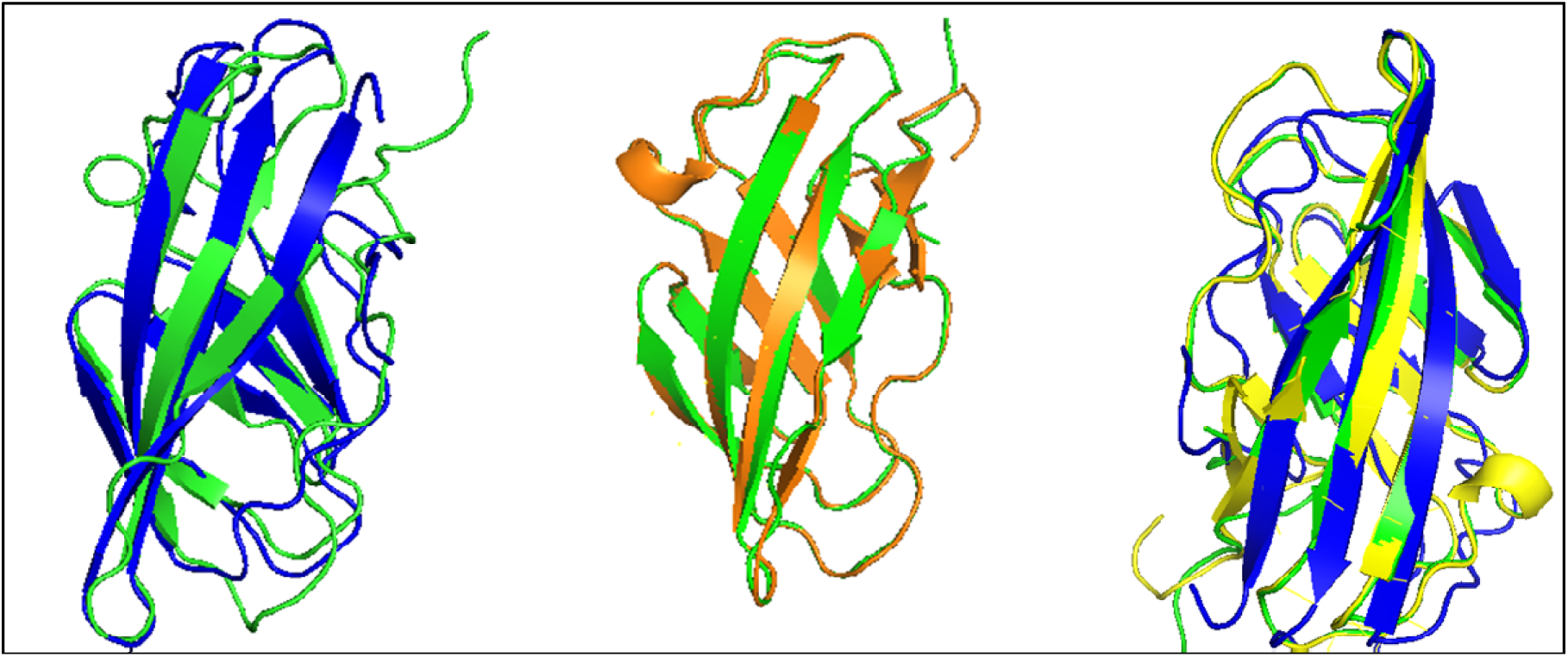
Structure based alignment of Rv1987 with chitin binding protein and cellulose binding protein. (i) Alignment of Rv1987 (green)with chitin-binding domain of *Streptomyces coelicolor* (PDB ID: 2rtt) (blue); (ii) Alignment of Rv1987 (green) with cellulose binding domain of *Cellulomonas fimi* (PDB ID:1exg) (orange); (iii) Superimposed structures of Rv1987 with the chitin binding domain/cellulose binding domain in (i) and (ii).

**Table 1.**
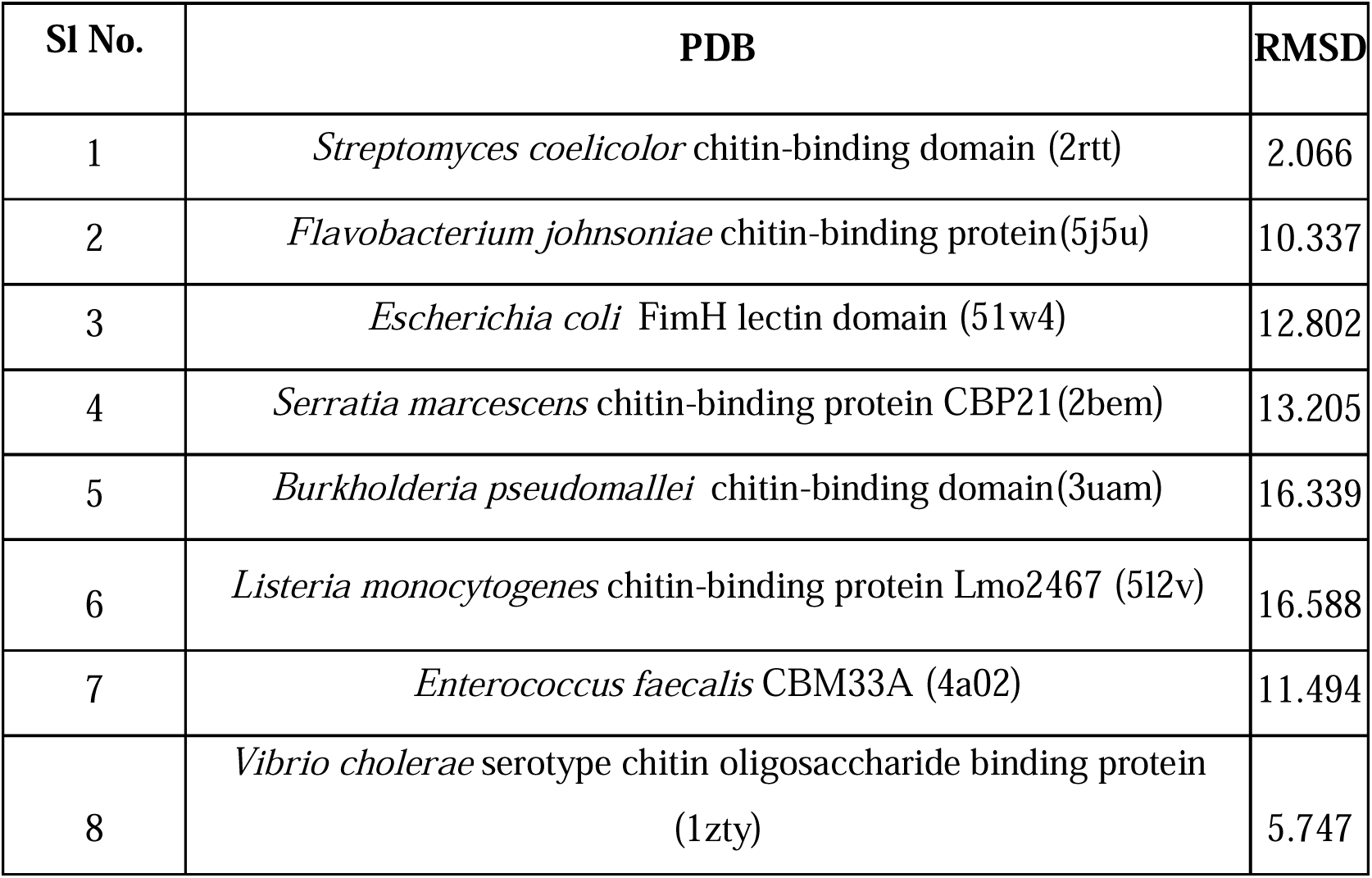
RMSD values from structural overlap analyses of chitin binding proteins with Rv1987.

**Table 2.** RMSD values from structural overlap analyses of cellulose binding proteins with Rv1987.

| SI No. | PDB | RMSD |
| --- | --- | --- |
| 1 | *Cellulomonas fimi* cellulose binding domain(1exg) | 0.289 |
| 2 | *Clostridium thermocellum* cohesin-2 domain (1anu) | 4.741 |
| 3 | *Cellvibrio japonicus* Type X Cbd (1e8r) | 6.846 |
| 4 | *Hypocrea jecorina* cellulose-binding domains (1azj) | 9.815 |
| 5 | *Trichoderma reesei* cellulose-binding domain (4bmf) | 10.132 |
| 6 | *Clostridium thermocellum* cellulose-binding domain(1nbc) | 12.767 |
| 7 | *Spirochaeta thermophila* - CBM64(5e9p) | 13.312 |
| 8 | *Clostridium josui* carbohydrate-binding module(3acf) | 15.635 |

Thus, the above-mentioned results conclude that structurally Rv1987 is highly identical to cellulose binding proteins followed by chitin binding proteins. Also, structurally Rv1987 is different from other bacterial chitinases characterized so far. These observations indicate that Rv1987 is a cellulose/chitin binding protein.

### Binding pockets of Rv1987 recognize chitin/cellulose polymers

Results from sequence-based and structure-based studies indicated that Rv1987 probably can be a cellulose/chitin binding protein rather than an enzyme. Therefore, flexible molecular docking of Rv1987 was carried out with 14 different carbohydrate substrates chosen based on similar relevant chemical nature and molecular weight. Though a model structure of Rv1987 has been used for these studies due to the non-availability of X-ray diffraction solved crystal structures. However, we have made all efforts towards the quality assessment of the used structure from structural genomics consortium database of *M.tuberculosis.* This structure shows more than 75% of model coverage for the Rv1987 with the template structure. Also, a very high match (0.2 RMSD) of Rv1987 structure with experimentally solved crystal structure of *C. fimi* (1EXG) cellulase as discussed above is another indicative of the structure quality.

Nine successful protein-substrate target complexes were generated per substrate. The structure complexes with the highest ΔG value was considered as the best orientation implicating protein ligand binding. As shown in Table 3, chitin octamer, NAGOS-6 and cellotetroase were the best substrates as indicated by their binding scores. The docked orientations were studied further using Protein-Ligand Interaction Profiler which plots 3D orientations and list outs interacting residues with the substrates. The interacting residues and 3D map are depicted in figure 8. Hydrophilic amino acids such as Ser, Thr and Asp were found to be the key residues involved in the formation of hydrogen bonds with the substrates thus contributing to higher binding affinity. High docking scores with the tested carbohydrate substrates indicate that Rv1987 exhibits a strong binding to cellulose and chitin substrates (Table 3).

**Figure 8.**
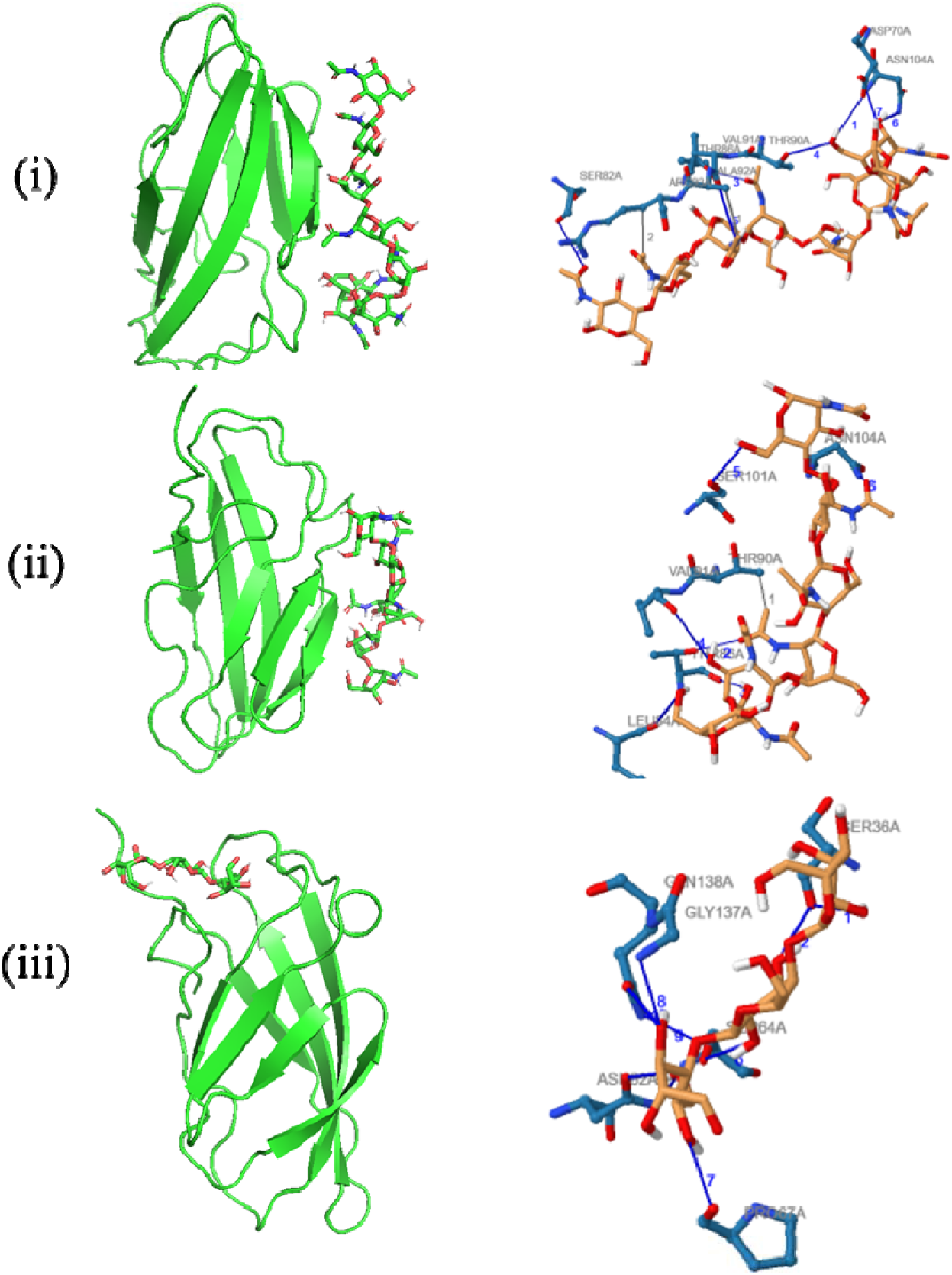
Analysis of binding pockets of Rv1987 to assess substrate specificity. 3D and 2D maps from molecular docking studies of different substrates with Rv1987 show (i) Chitin Octamer (ii) NAGOS-6 (iii) Cellotetraose as substrates exhibiting prominent interactions and high affinity with the binding pockets.

**Table 3.** Results of docking studies of Rv1987 with plausible carbohydrate substrates along with the binding energy values.

| **Ligands** | **Binding energy (kcal/mol)** |
| --- | --- |
| Chitin Octamer | -8.9 |
| NAGOS-6 | -8.9 |
| Cellotetraose | -8.3 |
| Core oligosaccharide | -8.1 |
| Triacetyl chitotriose | -7.6 |

### Novel Chitin/Cellulose-binding-domain protein Rv1987 in M.tuberculosis is cell wall/membrane localized

Rv1987 harbours a signal peptide and is also found to be present in the culture filtrate [32], a property not directly expected of entities that facilitate cellular interaction or adhesins. However, a proportion of such molecules might stick to the outer membrane of bacteria and mediate attachment to relevant substrates. *Bordetella pertussis* has been shown to secrete hemagglutinins containing carbohydrate recognition domains, which are involved in the adherence of bacteria to epithelial cells by binding to glycoconjugates on cell membranes [33], [34]. Another hemagglutinin of *V. cholerae* has been reported to be specific for N-acetyl-D-glucosamine, the monomeric subunit of chitin, and mediating binding of the bacterial cells to rabbit intestinal epithelial cells [35]. Furthermore, it has been shown that *Klebsiella pneumoniae* is internalized efficiently by cultured human epithelial cells, presumably *via* an N-glycosylated receptor containing N-acetyl-glucosamine residues [36].

Western blotting of whole cell lysates of *M. tuberculosis* strain H37Rv using anti-Rv1987 antibodies shows prominent bands at 14kDa (Figure 9). Presence of Rv1987 was detected in the culture filtrate, cell wall, cell membrane and in the whole cell lysates of virulent *M. tuberculosis*. Thus, presence of this protein in all the cell fractions indicates its wider distribution within the bacteria. The presence of Rv1987 in the cell wall fraction imply that the protein could also possibly be a structural component of the cell wall. While the presence in culture filtrate fraction suggests the secretory characteristic of Rv1987. There was also a prominent signal in the cell membrane fraction. Mycobacterial cell envelope consists of plasma membrane, arabinogalactan/peptidoglycan core which is cell wall and an outer membrane. In the experiment, the cell membrane preparation used (BEI resources-NR-14831) is a preparation of the cell membrane fraction that contains the cytoplasmic membrane and components of the outer membrane layer. These observations indicate presence of Rv1987 in the exterior most cell structures of *M. tuberculosis* and thus its potential to interact with the host cells.

**Figure 9.**
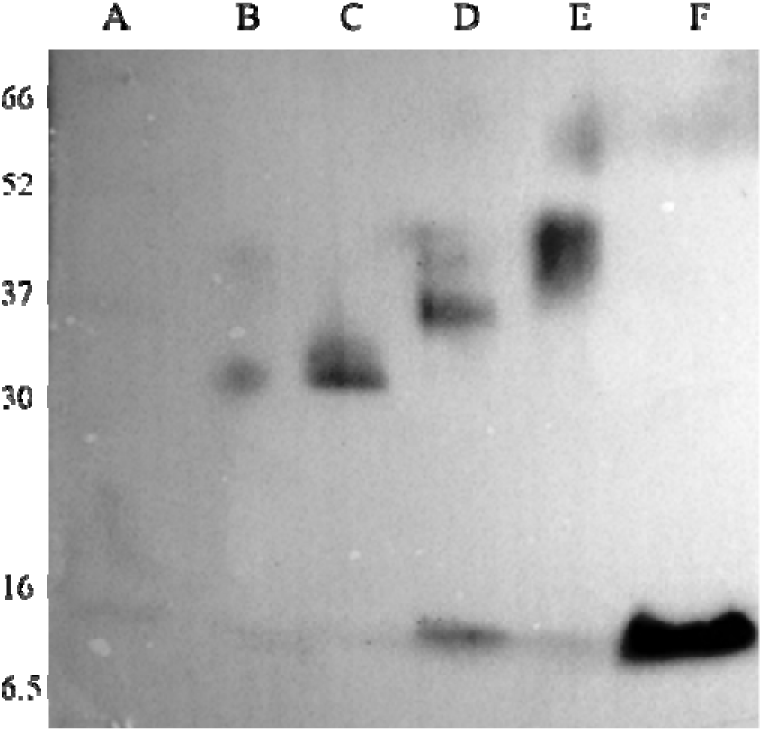
Analysis of sub-cellular localization of Rv1987 in *M. tuberculosis*. Detection of expression of Rv1987 in lysates of *M. tuberculosis* was carried out by western blotting using polyclonal mono specific anti-Rv1987 antibodies. Lane A: Molecular weight marker (kDa); Lane B: Soluble cell wall proteins; Lane C: Culture filtrate proteins; Lane D: Cell membrane fraction; Lane E: Whole cell lysate; Lane F: Purified protein Rv1987.

Speculatively, a glycosylated epithelial cell surface molecule might exist as receptor for Rv1987. These data suggest a role for Rv1987 in pathogenicity, for instance, as an adhesin mediating colonization of eukaryotic cells. However, the results suggest that functional role of Rv1987 in pathogenesis is most likely during the initial events of infection and might also play a role in Mycobacterial biofilm formation. It is important note that very recent reports have shown that *M. tuberculosis* form biofilms *in vivo* and these films are rich in cellulose [27] [28].

### Rv1987 influences bacterial interaction with the host cells

*M. tuberculosis* expresses adhesion proteins on its surface which aid in bacterial attachment to the host cell receptors during colonization. Adherence of *M. tuberculosis* to respiratory epithelium induces membrane perturbation and the formation of membrane extensions enable bacterial adherence to respiratory epithelium[37]. As the alveolar epithelial cells are the first cells of contact with the bacteria during infection in humans and monocytes/macrophages are the resident cells for Mycobacteria, we assessed interactions of recombinant *M. smegmatis* over expressing Rv1987 with these two cell types employing respective cell lines. To determine whether Rv1987 has any role in initial attachment to the host cells, thereby aiding the bacterial infection, recombinant *M. smegmatis*::Rv1987 was added to A549 lung epithelial cells for a brief period of 60, 120 and 180 minutes at a MOI (multiplicity of infection) of 10. Cell attached bacteria were enumerated through CFUs. *M.smegmatis*::Rv1987 had an enhanced adhesion effect in comparison to *M*.*smegmatis*:pVV16 (Figure 10a). These results indicate that Rv1987 could aid *M*. *tuberculosis* during the initial stages of infection.

**Figure 10.**
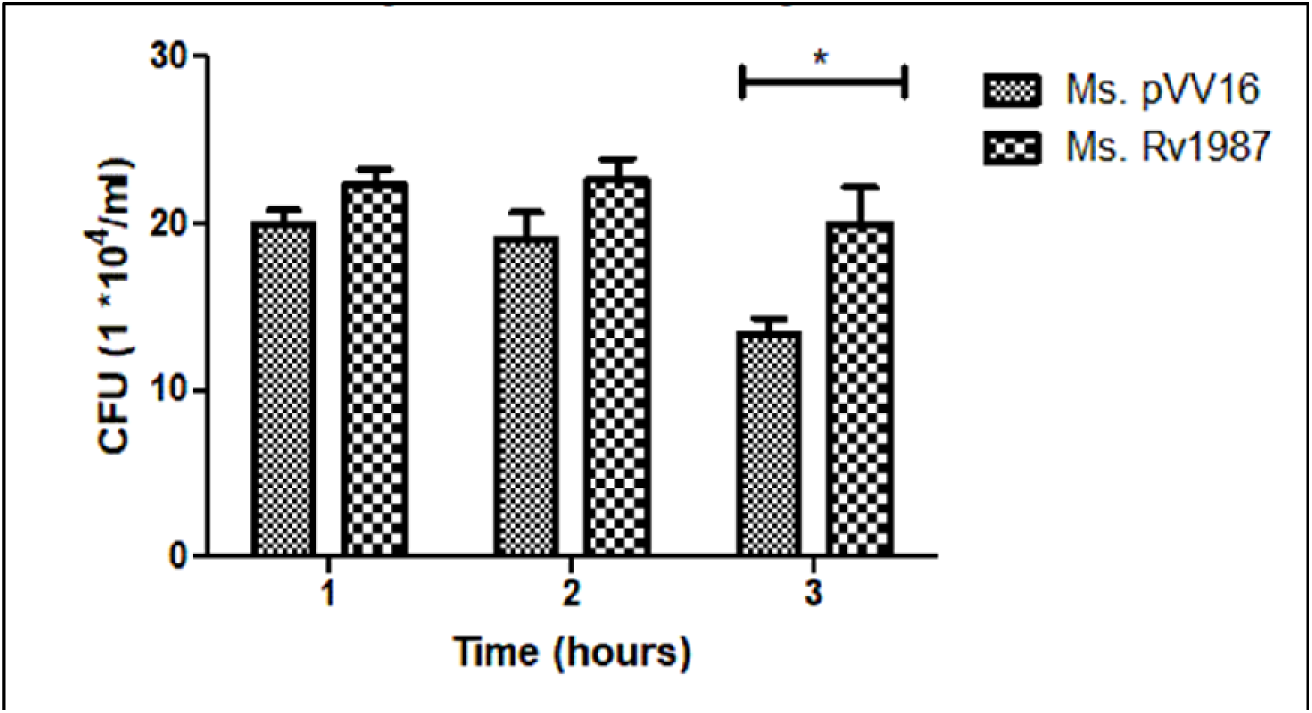
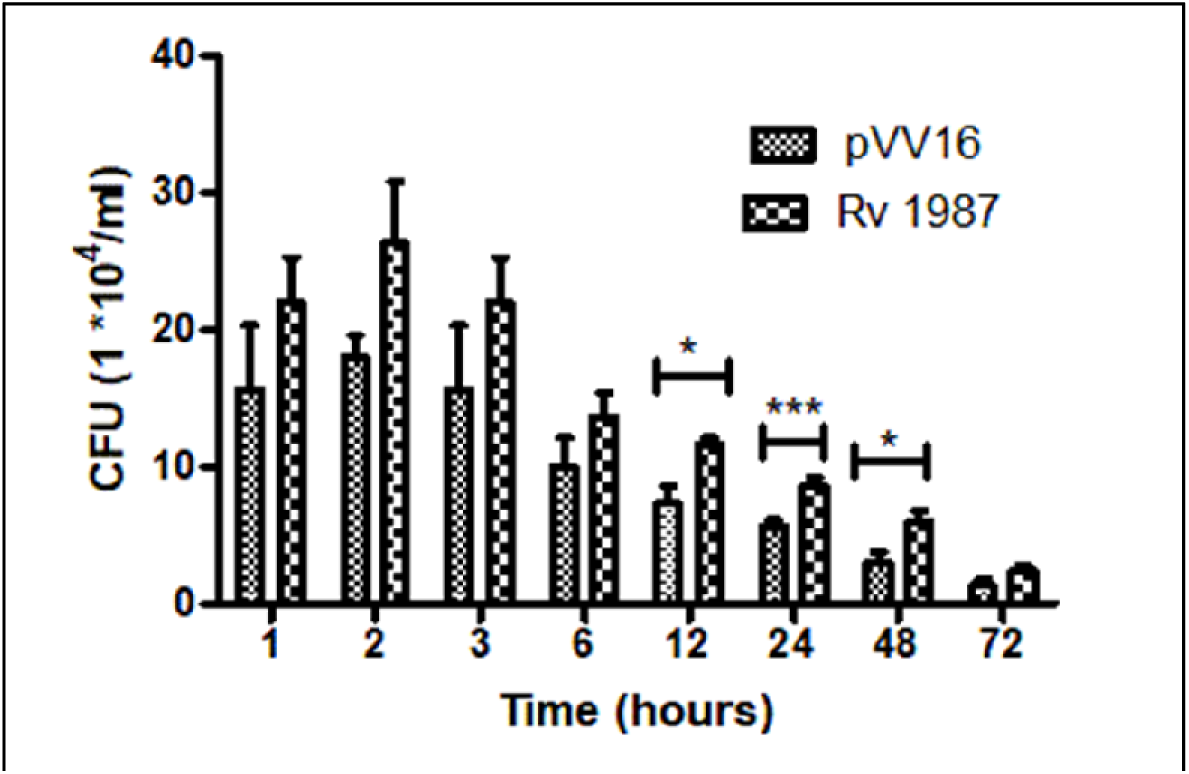
**a. Cytoadhesion assay.** Recombinant *M.smegmatis* harboring Rv1987 shows enhanced cytoadhesion to human lung epithelial cell line A549. Error bars represent standard deviation. *M.smegmatis* transfected with vector alone was used as a negative control. Statistical analyses were performed using graph pad prism 5 software. One way ANOVA analysis was used to determine the statistical difference between these two groups which is defined as follows: *p < 0.05. **b. Intracellular survival assay**. Recombinant *M. smegmatis* harboring Rv1987 shows enhanced survival in differentiated THP-1 cell lines post infection at 10 MOI. Error bars represent standard deviation. *M.smegmatis* transfected with vector alone was used as a negative control. Statistical analyses were performed using graph pad prism 5 software. One way ANOVA analysis was used to determine the statistical difference between these two groups which is defined as follows: *p < 0.05 and ***p<0.005.

Macrophages are the primary host cells of *M. tuberculosis* in the host post-infection and hence the survival within them is critical to determine the fate of the infection. In order to assess whether Rv1987 facilitates the intracellular survival of bacteria in macrophages, assays were carried out by infecting differentiated THP-1 cells with *M.smegmatis*::pVV16 and *M.smegmatis*::Rv1987 at an MOI of 10 for different late time points post infection. It was observed that the recombinant strain expressing Rv1987 showed almost 2-fold higher bacillary load in THP-1 macrophages at the end of 8 hours (figure 10b). These results suggest that Rv1987 is able to enhance intracellular survival of *M. smegmatis* within macrophages.

## Discussion

Worldwide burden of TB warrants aggressive efforts to improve the knowledge of the pathogen, the host, and their interactions so as to intervene with and reverse the progressive worsening of pulmonary and extra pulmonary TB. In this context, the present work addresses how a secreted cellulose and chitin binding protein which is normally thought to provide nutritional benefits for environmental bacteria in their natural habitats such as soil could be used by *M. tuberculosis*, a human pathogen.

Bacterial cellulose and cellulose binding proteins are only beginning to be characterized. It is important to note that several human pathogenic bacteria form biofilms rich in cellulose. Interestingly, recent research in animal models has shown that Mtb forms cellulose rich biofilms *in vivo* in tissue spaces and extra cellular matrix [27] [28] that could be allowing the bacteria to persist in the host. However, information on cellulose synthesis cluster in Mycobacteria is very sparse. Understanding biofilm dynamics formed *in vivo* is of clinical significance which will serve to therapeutically target these infections. Therefore, unravelling entities that could be stabilizing biofilms via the cellulose binding functions such as the likes of Rv1987 aptly fits the quests in the new dimension of TB research. Furthermore, bacterial adhesins are known to play an important role in the formation of biofilms. The observation that Rv1987 could be acting as adhesin as well, reiterates its possible role in stabilizing biofilms. Also, given that different bacteria synthesize completely different types of cellulose, it would be interesting to further probe into if the cellulose synthesis clusters and CBPs differ too.

Novel observation from our work is that Mycobacterial proteins presently annotated as chitinase(s) unlike their counterparts from other bacteria are unique in the sense that they show strong binding preference to cellulose too.

It is of interest to note that several components involved in mammalian immune responses are glycosylated and contain N-acetyl glucosamine residues linked by β1,4-glycosidic bonds and hence could be the targets for Rv1987. The present observation hints on possible ways by which pathogenic bacteria utilize carbohydrate binding proteins to interact with host cell components. To summarize, immediate outcome of the current study is an extended understanding of the host-pathogen interaction with respect to individual components of *M. tuberculosis.* Such studies will facilitate unravelling novel mechanisms of pathology which in turn will direct novel interventions to shorten the treatment course for TB.

## Acknowledgements

We thank Dr. Nagasuma Chandra, Department of Biochemistry, Indian Institute of Science, Bangalore, India, for the help with computational analysis. VJ acknowledges funding from SERB vide grant ECR/2015/000602. CMP acknowledges Indian council for medical research (ICMR), India for junior research fellowship. We acknowledge BEI resources, NIAID, NIH for the generous supply of cell fractions of *M. tuberculosis* strain H37Rv.

## Conflict of Interest statement

The authors declare that they have no competing interests.

## Author contribution

**CMP:** Performed the experiments, Validation, Data analysis, editing of manuscript draft final version; **VJ**: Conceptualization, Supervision, Data analysis, Original draft writing.

## Data availability statement

Experimental data that support the findings of this study are available in the results section of this article. Most of the software used for computational analysis described in materials and methods section are available on the public domain and have been referenced accordingly. Protein structures used are from protein data bank and mycobacterial structural genomics consortium and the relevant literature has been cited appropriately. The other raw datasets generated and/or analysed during the study are available from the corresponding author on reasonable request. We declare that we will be fully willing to comply with the journal policy and will be able to make any materials/data available required for the review process and thereafter.

## Notes

### Competing Interest Statement

The authors have declared no competing interest.

